# Experimental infection of *Artibeus lituratus* bats and no detection of Zika virus in neotropical bats from French Guyana, Peru, and Costa Rica, suggest a limited role of bats in Zika transmission

**DOI:** 10.1101/2022.04.25.489338

**Authors:** Alvaro Aguilar-Setién, Mónica Salas Rojas, Guillermo Gálvez Romero, Cenia Almazán Marín, Andrés Moreira Soto, Jorge Alfonso-Toledo, Cirani Obregón Moralesn, Martha García Flores, Anahí García Baltazar, Jordi Serra-Cobo, Marc López-Roig, Nora Reyes Puma, Marta Piche-Ovares, Mario Romero-Vega, Daniel Felipe Barrantes Murillo, Claudio Soto-Garita, Alejandro Alfaro Alarcón, Eugenia Corrales-Aguilar, Osvaldo López-Díaz, Felix Drexler

**Author notes:** **Author Contributions** Conceptualization: Álvaro Aguilar Setién, Jean Félix Drexler, Formal analysis: Cenia Almazán Marín, Mónica Salas Rojas, Guillermo Gálvez Romero, Jorge Alfonso Toledo, Martha García Flores, Andrés Moreira Soto. Funding acquisition: Alvaro Aguilar Setién and Jean Felix Drexler. Investigation: Cenia Almazán Marín, Mónica Salas Rojas, Gillermo Galvez Romero, Jorge Alfonso Toledo, Martha García Flores, Anahí García Baltazar, Cirani Obregón Morales, Andrés Moreira, Methodology: Cenia Almazán Marín, Mónica Salas Rojas, Guillermo Gálvez Romero, Jorge Alfonso Toledo, Martha García Flores, Anahí García Baltazar, Cirani Obregón Morales, Andrés Moreira, Jordi Serra-Cobo, Marc López-Roig, Nora Reyes Puma, Marta Piche-Ovares, Mario Romero-Vega, Daniel Felipe Barrantes Murillo, Claudio Soto-Garita, Alejando Alfaro-Alarcón, Osvaldo Lopez-Díaz, Eugenia Corrales-Aguilar. Project administration: Alvaro Aguilar Setién and Jean Felix Drexler. Resources: Alvaro Aguilar Setién and Jean Felix Drexler. Supervision: Alvaro Aguilar Setién and Jean Felix Drexler. Visualization: Cenia Almazán Marín, Mónica Salas Rojas, Martha García Flores, Alvaro Aguilar Setién Writing – original draft: Alvaro Aguilar Setién, Jean Felix Drexler, Cenia Almazán Marín, Guillermo Gálvez Romero, Andrés Moreria Soto. Writing – review & editing: Alvaro Aguilar Setién, Jean Felix Drexler, Andrés Moreria Soto, Jordi Serra-Cobo.

## Abstract

Bats are important natural reservoir hosts of a diverse range of viruses that can be transmitted to humans and have been suggested that could play an important role in the Zika virus (ZIKV) transmission cycle. However, the exact role of these animals as reservoirs for Flaviviruses is still controversial. To further expand our understanding of the role of bats in the ZIKV transmission cycle in Latin America, we carried an experimental infection in wild-caught *Artibeus* bats and sampled several free-living neotropical bats over three countries of the region. Experimental ZIKV infection was made in free-ranging adult bats (4 females and 5 males). The most relevant gross findings were hemorrhages in the bladder, stomach and patagium. Significant histological findings included inflammatory infiltrate consisting of a predominance of neutrophils and lymphocytes, in addition to degeneration in the reproductive tract of males and females. This suggests that bat reproduction might be at some level affected by ZIKV. Leukopenia was also observed in some inoculated animals. Hemorrhages, genital alterations, and leukopenia are suggestive to be caused by ZIKV, however, since these are wild-caught bats, we can not exclude other agents. Excretion of ZIKV by qPCR was detected (low titles) in only two urine samples in two inoculated animals. All other animals and tissues tested negative. Finally, no virus-neutralizing Abs were found in any animal. To determine ZIKV infection in nature, a total of 2056 bats were blood sampled for ZIKV detection by qPCR. Most of the sampled individuals belonged to the genus *Pteronotus* sp. (23%), followed by the species *Carollia* sp. (17%); *Anoura* sp. (14%), and *Molossus* sp. (13.7 %). No sample of any tested species resulted positive to ZIKV by qPCR.

These results together suggest that bats are not efficient amplifiers or reservoirs of ZIKV and may not have an important role in ZIKV transmission dynamics.

**Author summary:** In previous works made in 2008-2009, we have found the presence of antibodies against Flaviviruses and viral RNA has been detected in Neotropical chiropterans of Mexico, which led us to support the hypothesis that these animals could be reservoirs of Flaviviruses. As controversial opinions have been exposed, and based on a previous (2019) experimental ZIKV infection made in Colorado State University using adult *Artibeus* males from a captive colony, in this work we also experimentally infected adult *Artibeus* males complementarily adding females and using free-living animals instead of laboratory bats. We also monitored a diverse range of natural bat populations in Latin America for the presence of viral RNA against ZIKV in blood. A plaque reduction seroneutralization test was used for the detection of antibodies against ZIKV. Similar to the previous work, we found histopathological alteration in male testicles but also in ovaries and oviducts of females, as well as gliosis and multifocal necrosis in pyramidal neurons and Purkinge cells of inoculated animals. Only two urine samples from inoculated animals showed viral RNA. Additionally, leukopenia and lymphoid follicular splenic hyperplasia were evidenced. Differing to what was reported, no neutralizing antibodies against ZIKV were detected in any sample. Viral RNA within the blood was not present in any of the 2056 bat samples collected in French Guyana, Peru and Costa Rica and proceeding from 33 bat genera. These results together suggest that bats are not efficient amplifiers or reservoirs of ZIKV and might not have an important role on ZIKV transmission dynamics.

## Introduction

Zika virus (ZIKV) belongs to the genus *Flavivirus* inside the Flaviviridae family (1). ZIKV was first isolated in Africa in 1954 (1, 2). The virus circulates between an urban transmission cycle involving arthropod vectors and humans, and a sylvatic cycle involving arthropod vectors and non-human primates (2, 3). In addition, ZIKV alternative transmission routes in humans include sexual, blood transfusion and perinatal (4-7). ZIKV was first reported in the Americas in 2015 and since then it spread throughout the continent causing more than 850,000 human cases (8). During the outbreak, serological surveys performed in Brazil detected a high 60% population exposure(8). This high population exposure suggested the end of the outbreak due to a high herd immunity. However, if ZIKV adapts to new vertebrate wildlife hosts in the Americas like Yellow fever virus (YFV) (9), ZIKV might stablish a sylvatic cycle until there is enough naïve population to cause another outbreak.

Early during the outbreak, modeling studies suggested non-human primates as the most probable ZIKV hosts (10). However, serological surveys in wild non-human primates in Brazil found a limited role in ZIKV transmission and urged to study other wildlife species as possible ZIKV hosts (11, 12). Considering the vertebrate species richness in America, the role of non-primate animals in the ZIKV cycle remains understudied (13). Bats are a megadiverse group of mammals only outnumbered by rodent diversity and in the American continent. Molecular and serological detection of flaviviruses suggest exposure to flaviviruses in the wild (14-19). No active or transmissible flavivirus infection in wild bats has been reported. For the most studied flavivirus in bats, DENV, a peri-domestic study found only limited exposure of bats likely due to closeness to humans and consumption of DENV vectors (20). Additionally, *in-vitro* studies infecting several neotropical bat cell lines, searching serological and molecular evidence of infection in wild bats, and experimentally infecting *Artibeus* bats with Dengue virus (DENV) showed that these species are inadequate DENV hosts and may do not play an important role in DENV transmission (21-24). For ZIKV, anecdotal experimental infections and field studies performed in the 50s-60s have documented the susceptibility and disease development in African and American bat species (25-27). Recently, an experimental infection in a breeding colony of *Artibeus jamaicensis* bats detected ZIKV RNA and seroconversion in some studied animals, and raised the possibility that bats may have a role in Zika virus ecology even endangering bat populations (28). Considering the neotropical bat species richness and limited information of ZIKV hosts to date, we carried an experimental infection in wild-caught *Artibeus* bats and sampled free-living neotropical bats over several Latin American countries to assess their role in the ZIKV transmission cycle in the Americas.

## MATERIALS AND METHODS

### Capture of bats for infection experiments

Eleven great fruit-eating bats (*Artibeus lituratus*) were captured in Oaxtepec, Mexico. Animals were measured, weighed, aged, sexed, tagged and identified as *Artibeus lituratus* species using a dichotomous key by field specialists. Capture and animal handling was approved by the Mexican environmental standards (permit SEMARNAT Mexico No SGPA/DGVS/08986/18). To ensure that the animals were not acutely infected by ZIKV, urine and plasma samples were taken after capture and tested with a ZIKV-specific qRT-PCR (29). Next, bats were quarantined for 1 month before the experiment. Nine bats were selected for the experimental infection and two as controls. Animals were kept in captivity with food and water *ad-libitum*, according to the Guidelines of the American Society of Mammalogists for the Use of Wild Mammals in Research (30).

### Experimental infection

#### Virus stock

A ZIKV isolate from a human patient in Yucatan, Mexico (ZIKV/Mer.IPN01) in 2017 was used. Briefly, C636 cells previously grown with 12 ml of Eagle’s minimal essential medium (DMEM) with 10% fetal bovine serum (FBS) (GIBCO®) were inoculated with ZIKV at a multiplicity of infection (MOI) of 0.1, incubated in DMEM without FBS at 27 °C for 1 hour after which new medium with FBS was added. The infected cells were kept until cytopathic effect was observed in >80% of the cells. Next, cell supernatant was collected, centrifuged and stored in aliquots at −70 °C. The viral stock titers were determined in BHK-21 cells using standard plaque assays in carboxymethylcellulose (CMC).

#### Experimental infection

Nine bats were injected subcutaneously with 2×10^5^ plaque-forming units (PFU) of Zika virus and two control bats with PBS in the scapular area the same day (**Table 1A)**. To observe ZIKV pathogenesis over time, bats were divided into 4 groups by sex: group 1 consisted of male MO04 and female MO06 euthanized 3 days post-inoculation (d.p.i.), group 2 male MO01 and female MO08 euthanized 7 d.p.i., group 3 males MO05 and MO02 and female MO09 euthanized 14 d.p.i. and group 4 male MO07 and female MO10 euthanized 21 d.p.i. Euthanasia of the control group of female MO11 and male MO12 was performed 21 d.p.i. (**Table 1 A)**.

**Table 1.**
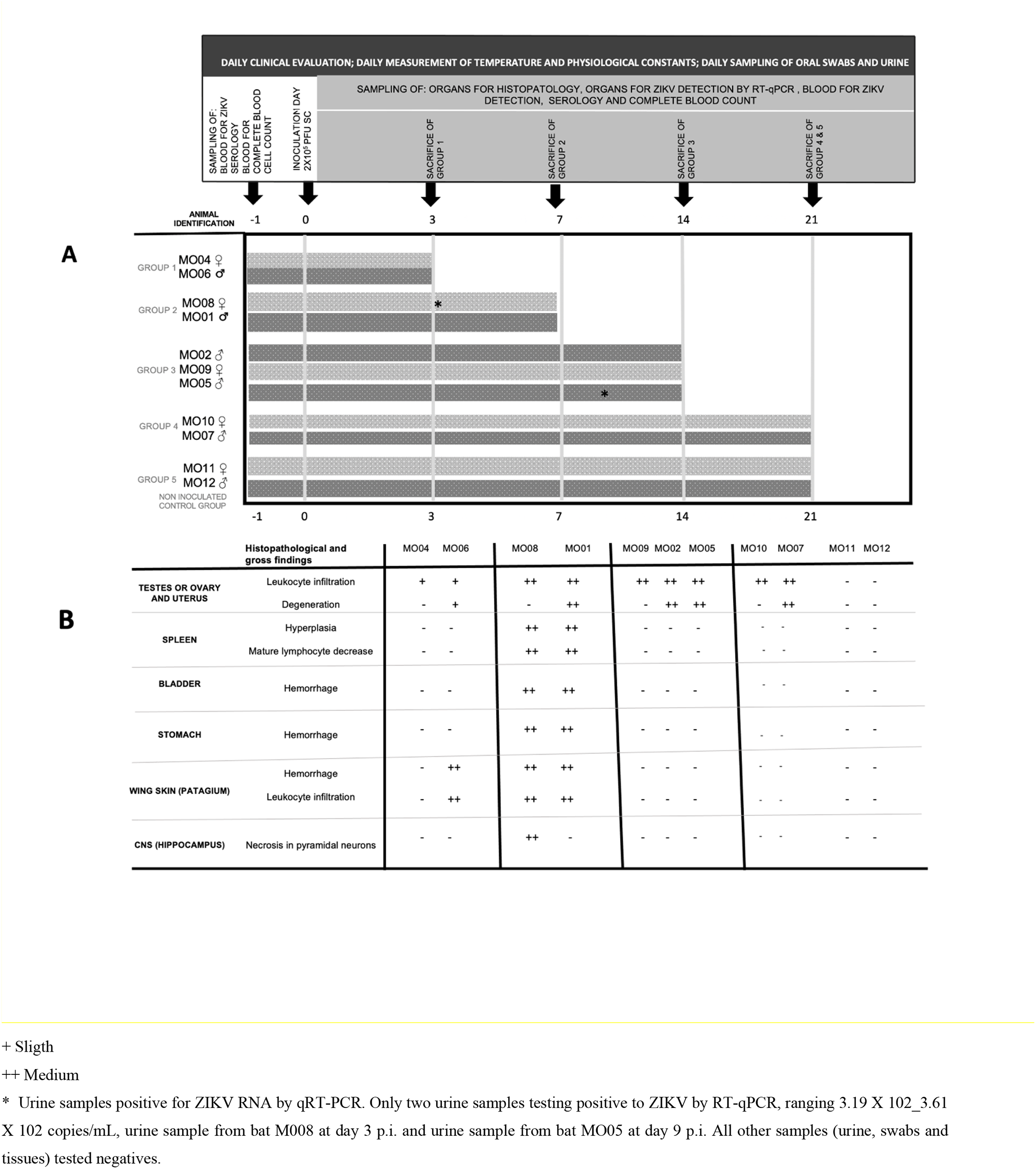
Experimental design and timeline. Bats were inoculated (SC) with 2 × 10^5^ UFP of ZIKV Mer.IPN01. Groups, sampling, and time of sacrifice A. Histopathological and gross findings B.

Bats were observed daily, body temperature and weight were recorded. Behavioral changes or signs suggestive of diseases such as lethargy, nasal or oral discharge, urine color, and pasty and or diarrheic feces consistency were recorded. Oral swabs and urine were collected daily (**Table 1A)**. On the day of euthanasia, bats were anesthetized using 30 μl a mixture of 1:1 of Ketamine 100 mg/mL–Xylazine 100 mg/mL and euthanized by cardiac exsanguination. Next, blood samples were divided for molecular and serological ZIKV detection and whole blood for complete blood count (CBC). The CBC was made in a VetScan HM5 device previously calibrated for Phyllostomid bat leucocytes using the reference previous reports (31-34). Differential counts WBC were made also in blood smears stained with Giemsa dye following standard protocols. Different organs such as brain, tongue, salivary glands, heart, lung, liver, spleen, gut, kidney, bladder, uterus, ovary, and testicles were collected on the day of euthanasia of each group. Each sampled organ was divided into two parts. A part of each organ was stored at −80 °C until processed for ZIKV-RNA detection by qRT-PCR. The remaining part of each organ was fixed in 10% buffered formalin and routinely processed for histopathological evaluation.

### Laboratory analyses

#### Histological studies

The formalin and paraformaldehyde-fixed tissues were processed using standard methodology: paraffin-embedded, 5 μm thickness sectioned and stained with Hematoxylin and Eosin (H&E).

#### Plaque reduction neutralization test

Plaque reduction neutralization test (PRNT) using ZIKV strain H/PF/2013 was performed Briefly, sera were diluted 1:10, 1:40, 1:100 and 1:350 in serum-free DMEM. Subsequently, a mixture of 35 μL of each serum dilution plus 35 μL of ZIKV was incubated for 1 hour at 37 ° C with 5% CO_2_. After this time, 50 μL of the serum dilution and ZIKV mixture plus 250 μL of cell culture media was added to previously seeded Vero cells and incubated for 1 hour at 37 ° C with 5% CO_2_. After the incubation cell media was changed and cells were incubated for 4 days, fixed with 6% paraformaldehyde and plaques visualized using violet crystal solution. A positive result was measured when a serum dilution reached < 50% of the number of plaques counted in the controls (35).

#### Real-Time RT-PCR

RNA extraction for the experimental infection and the sampling was carried out using MagNA Pure 96 DNA and Viral NA Small Volume Kit, in the MagNA pure 96 system (Roche, Germany). The presence of ZIKV was detected using a ZIKV-specific real-time PCR described previously (29).

### Fieldwork

#### A sampling of bats in different countries of Latin America

All sampling and capture were done with the proper wildlife permits and ethics clearances of the host countries-Samples were gathered before and after the ZIKV epidemics in the Americas between 2010 and 2019 in Peru, Costa Rica, and French Guyana. The complete data including sampling sites and geographic coordinates are included in the Table S1. Bats were captured using mist nets in neotropical forests, degraded forests, urban sites, and caves not targeting any specific bat species. After the capture, bats were kept individually in cotton bags until examination. Trained specialists identified the species using the morphological keys to the species. Blood samples for ZIKV detection by qPCR were taken by a trained veterinarian either from the brachial vein in case the animal was being marked and released, or intracardiac puncture after intramuscular application of euthanasia.

## RESULTS

### Experimental infection

#### Clinical signs of ZIKV disease in bats

None of the captured bats was ZIKV positive though qPCR before the experiment. For the nine adult *Artibeus lituratus* bats and the two controls, no clinical signs suggestive of disease were recorded during the quarantine period. The inoculated bats showed a mean temperature (geometric mean: 35.6 ºC; SD 0.75; CI 95%), similar to the non-infected control bats (geometric mean: 35.7 ºC; SD 0.6; CI 95%). Two inoculated bats showed extreme temperatures (male MO07, at day 3 p.i. 38.5 ºC, and male MO02 at day 6 p.i. 38.3 ºC) (Figure S1); this temperature is not at odds with the mean temperature for Phyllostomid bats that ranges between 34.5 and 39.5 ºC (36, 37), a suggesting that bats do not show fever when infected or other physiological alterations, as observed for several other viruses found in bats as Coronavirus (38, 39) and Marburg virus (40)

#### Complete blood count

The most relevant results in the red blood cell count show some bats with decreases in hemoglobin, in addition all individuals presented some low erythrocyte indices from day 0 to day 21 after inoculation, there being a clear normochromic microcytic anemia in M2 on day 14 post-inoculation and M9 presented a marginal erythrocytosis on day 21 postinoculation. The most relevant results in the white blood cell count compared to the values given for *Artibeus* bats (4.7 ± 2.4 × 10^9^/L) (31-34) (Table 2). show leukocytosis with lymphocytosis and monocytosis in individuals M6 and M8, in addition to lymphocytosis in M9 and leukocytosis with lymphocytosis in M11. Between 3 and 7 days after infection, a clear leukopenia with neutropenia was observed in the animals M4, M6, M1, M8, with recovery after day 9 pi. Individuals M2 and M9 presented neutropenia with lymphocytosis at 14 days pi where M9 showed a decrease in neutrophils with respect to day 0. The M5, M7, M10 and M11 only presented lymphocytosis.

**Table 2.** Complete blood count

| Parameters | References values | Post-infection days |  |  |  |  |  |  |  |  |  |  |  |  |
| --- | --- | --- | --- | --- | --- | --- | --- | --- | --- | --- | --- | --- | --- | --- |
|  |  | 0 |  |  | 3 |  | 7 |  | 14 |  |  | 21 |  |  |
|  |  | M6 | M8 | M9 | M4 | M6 | M1 | M8 | M5 | M2 | M9 | M7 | M10 | M11 |
| RBC 10<sup>12</sup>/L | 10.0-13.0 | 11.79 | 11.79 | 12.36 | IS | IS | IS | IS | 11.54 | 9.53 | 11.43 | 11.76 | 12.33 | 13.09 |
| HGB g/L | 155-195 | 156 | 154 | 174 | IS | IS | IS | IS | 153 | 132 | 139 | 148 | 155 | 159 |
| HCT L/L | 0.50-0.65 | 0.525 | 0.599 | 0.598 | IS | IS | IS | IS | 0.485 | 0.397 | 0.535 | 0.518 | 0.567 | 0.591 |
| MCV fL | 46.5-53.5 | 44.5 | 50.8 | 48.4 |  |  |  |  | 42.0 | 41.7 | 46.8 | 44.0 | 46.0 | 45.1 |
| MHC pg | 14.3-16.3 | 13.2 | 13.1 | 14.1 |  |  |  |  | 13.3 | 13.9 | 12.2 | 12.6 | 12.6 | 12.1 |
| MCHC g/L | 290-322 | 297.1 | 257.1 | 291 |  |  |  |  | 315.5 | 332.5 | 259.8 | 285.7 | 273.4 | 269 |
| WBC 10<sup>9</sup>/L | 2.3-7.1 | 7.62 | 8.26 | 6.88 | 0.7* | 0.3* | 0.5* | 0.9* | 3.64 | 5.78 | 2.58 | 5.43 | 6.50 | 8.13 |
| Neutrophils | 1599.35- 5420.81 | 3536 | 3643 | 3347 | 287 | 114 | 265 | 144 | 1762 | 1422 | 289 | 1695 | 2873 | 4057 |
| Band neutro. | 0.0-46.2 | 0 | 0 | 0 | 0 | 0 | 0 | 0 | 0 | 0 | 0 | 0 | 0 | 0 |
| Lymphocytes | 95.05 - 1570.01 | 3239 | 3866 | 3227 | 350 | 162 | 230 | 702 | 1594 | 4312 | 2118 | 3394 | 3471 | 3845 |
| Monocytes | 20.91 - 528.47 | 846 | 752 | 269 | 35 | 24 | 5 | 54 | 284 | 46 | 173 | 386 | 156 | 228 |
| Eosinophils | 0.0 -193.67 | 0 | 0 | 0 | 0.021 | 0 | 0 | 0 | 0 | 0 | 0 | 0 | 0 | 0 |
| Basophils | 0.0 -37.58 | 0 | 0 | 0 | 0.007 | 0 | 0 | 0 | 0 | 0 | 0 | 0 | 0 | 0 |
| Neutrophils% | 32.5-74.1 | 46.4 | 44.1 | 48.8 | 42.0* | 38.0* | 53.0* | 16.0* | 48.4 | 24.6 | 11.2 | 30.4 | 44.2 | 49.9 |
| Band % | 0-1 | 0 | 0 | 0 | 0* | 0* | 0* | 0* | 0 | 0 | 0 | 0 | 0 | 0 |
| Lymphocytes% | 2.0-33.6 | 42.5 | 46.8 | 46.9 | 50.0* | 54.0* | 46.0* | 78.0* | 43.8 | 74.6 | 82.1 | 62.5 | 53.4 | 47.3 |
| Monocytes % | 0.4-11.1 | 11.1 | 9.1 | 4.3 | 5.0* | 8.0* | 1.0* | 6.0* | 7.8 | 0.8 | 6.7 | 7.1 | 2.4 | 2.8 |
| Eosinophils % | 0-4.1 | 0 | 0 | 0 | 3* | 0* | 0* | 0* | 0 | 0 | 0 | 0 | 0 | 0 |
| Basophils % | 0-0.8 | 0 | 0 | 0 | 1* | 0* | 0* | 0* | 0 | 0 | 0 | 0 | 0 | 0 |

\* determination by blood smears assessment
IS, insufficient sample
Red letters, values below the reference; blue letters, values above the reference.
M, bat number

#### Gross findings

During the necropsy the main macroscopic lesions were observed in three bats, MO06 presented few petechial hemorrhages in the patagium 3 days p.i., MO01 and the MO08 presented mild to moderate petechial hemorrhages in the patagium and in the stomach mucosa and MO08 presented moderate petechial hemorrhages in the bladder mucosa that caused hematuria at 7 d.p.i. No further apparent pathological changes were observed in the necropsy; the macroscopic lesions found in infected bats suggest as a possible cause a ZIKV infection, however, since these are bats captured in the wild, we cannot exclude the presence of other agents such as filariasis (41) or DENV infection that cause bleeding in the skin and mucous membranes during experimental infections in *Artibeus* bats (23). In summary, because these lesions could occur in various physiopathogenies, more data are needed to consider them characteristic findings in ZIKV infections.

#### Histopathology

The males infected of the four groups demonstrated histological changes within the epididymis and testicle described as follows. The epididymal epithelium is markedly vacuolated, with loss of stereocilia (Figure 1 A-B). A mild interstitial lymphocyte infiltrates and mild to moderate germ cell degeneration were observed in the epididymis. Epididymal ducts are filled by degenerate to apoptotic spermatids. In the testicle, the seminiferous tubules have early stages of testicular degeneration, with marked vacuolization of the Sertoli cells, with evidence of apoptotic spermatids (Figure 1C-D) in addition to mild congestion and few to moderate neutrophils and macrophages infiltrating the interstitium.

**Figure 1.**
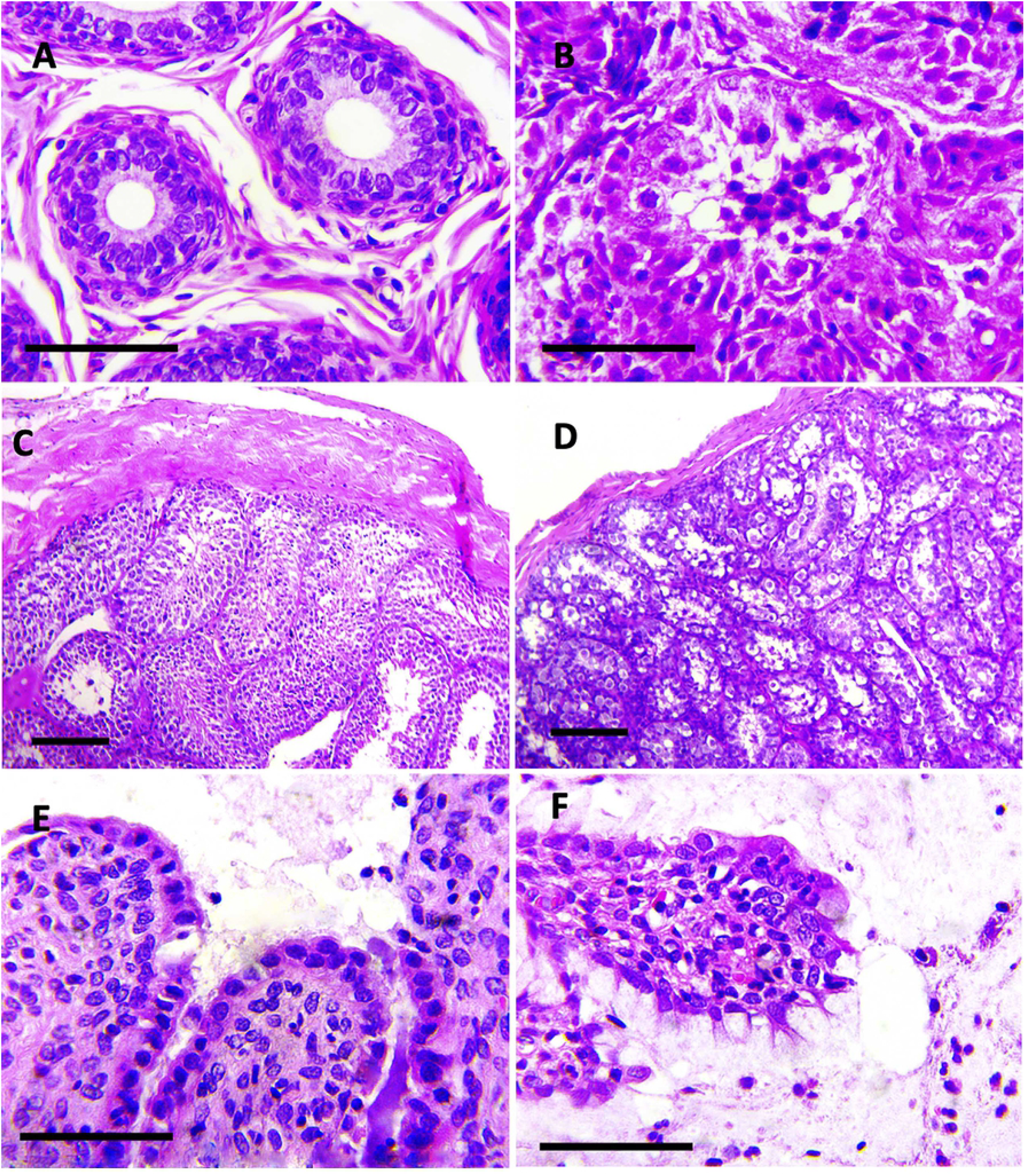
Photomicrographs of sections of the epididymis. A. Control epididymal ducts. H&E stain; bar = 50 μm. B. Epididymal duct epithelial degeneration, vacuolization, and loss of stereocilia, the lumen is filled by abundant degenerate to apoptotic spermatids. H&E stain; bar = 50 μm. Photomicrographs of sections of the testicles. C. Control testicular parenchyma. D. Early stage of testicular degeneration, vacuolated Sertoli cells with disorganized exfoliated of absent germ cells. H&E stain; bar = 50 μm. Photomicrographs of sections of the oviduct. E. The lamina propria is infiltrated by scant neutrophils. F. The degenerated epithelium has lymphocytic exocytosis and the oviduct lumen is filled by granular eosinophilic proteinaceous secretion intermixed with neutrophils. H&E stain; bar = 50 μm.

Similarly, all infected females presented mild to moderate lymphocytic infiltrates within the ovary and uterus throughout all the days after inoculation. The oviduct presented a minimal neutrophilic infiltrate within the lamina propria and degeneration of secretory and epithelial cells that was progressively increasing until day 21 without showing regression and lymphocytic exocytosis. The uterine lumen is filled by moderate amount of a granular eosinophilic proteinaceous secretion intermixed with a mild neutrophilic exudate (Figure 1 E and F).

The most representative lesions observed in the brain of the bat MO08 sacrificed 7 days after nfection, were present in the hippocampus; which showed areas with moderate multifocal necrosis in pyramidal neurons, in addition with moderate multifocal gliosis areas. Multifocal necrosis of the Purkinje cells was present within the cerebellum (Figure 2).

**Figure 2.**
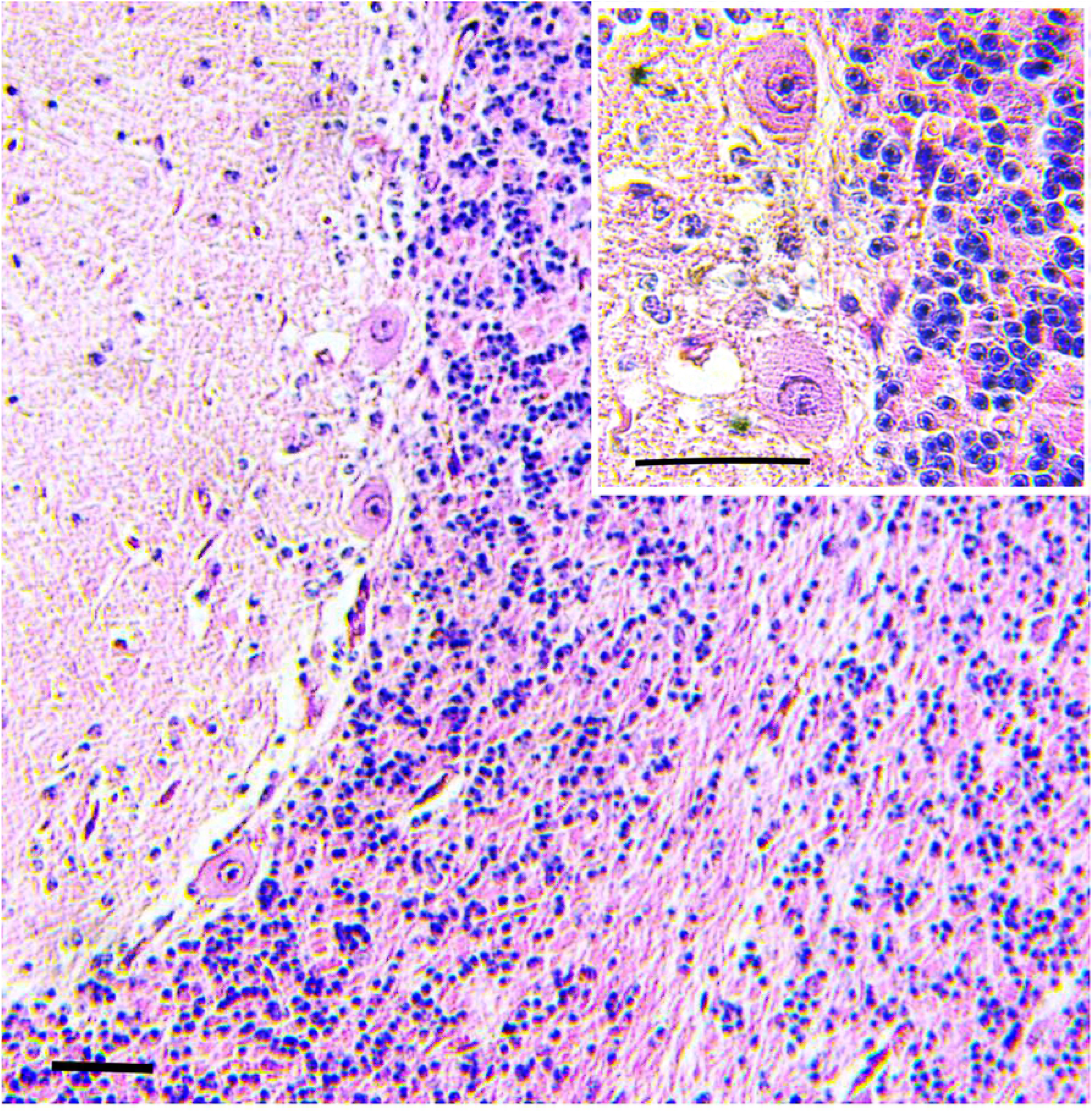
control. Photomicrographs of sections of the cerebellum of a control. Molecular, Purkinje cell, and granular layer. H&E stain; bar = 50 μm. Purkinje cells *(inset)*. H&E stain; bar = 50 μm.

**Figure 2.**
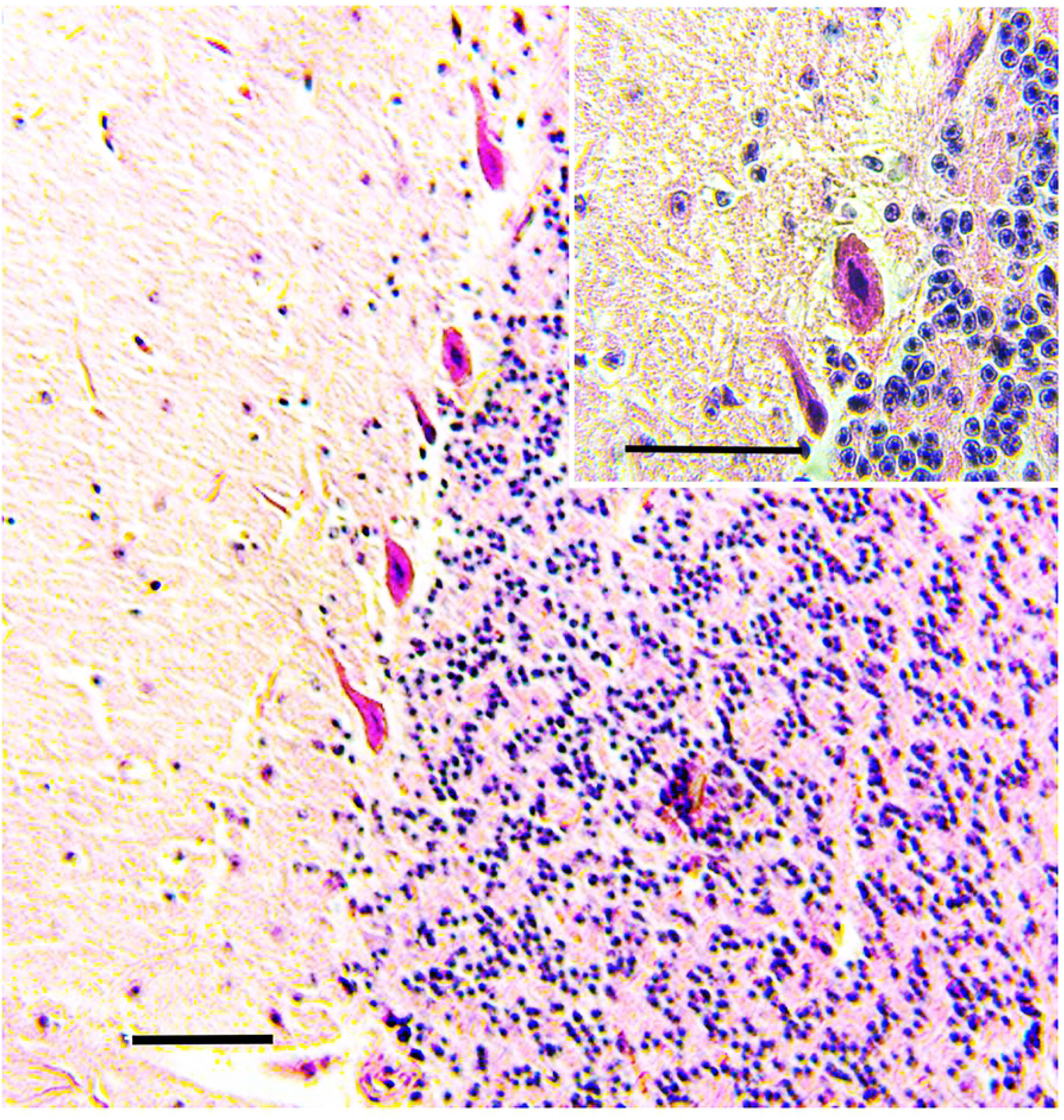
infected. Photomicrographs of sections of the cerebellum. Necrosis of Purkinje cell layer. H&E stain; bar = 100 μm. Necrotic cells are shrunken, with angulated cellular margins, hypereosinophilic cytoplasm, and pyknotic nuclei *(inset)*. H&E stain; bar = 50 μm.

The bats MO06, MO01 and MO08, presented moderate dermal hemorrhages in the patagium. The dermis was moderately infiltrated by neutrophils and lymphocytes (Figure 3). Dermal capillaries are markedly distended, congested with neutrophilic leukostasis. Finally, the spleen of MO01 and MO08, sacrificed 7 days pi (Group 2), showed moderate lymphoid depletion and follicular lymphoid hyperplasia (Figure 4). This observation is consistent with the WBC of these individuals who presented leukopenia due to neutropenia and additionally the total number of lymphocytes decreased drastically compared to the individuals sampled on day 0 including M008 which was sampled on day 0 and day 7 (Table 2). This finding is compatible with what has been reported during multiple viral infections that generate hypo-proliferation or lympholysis.

**Figure 3.**
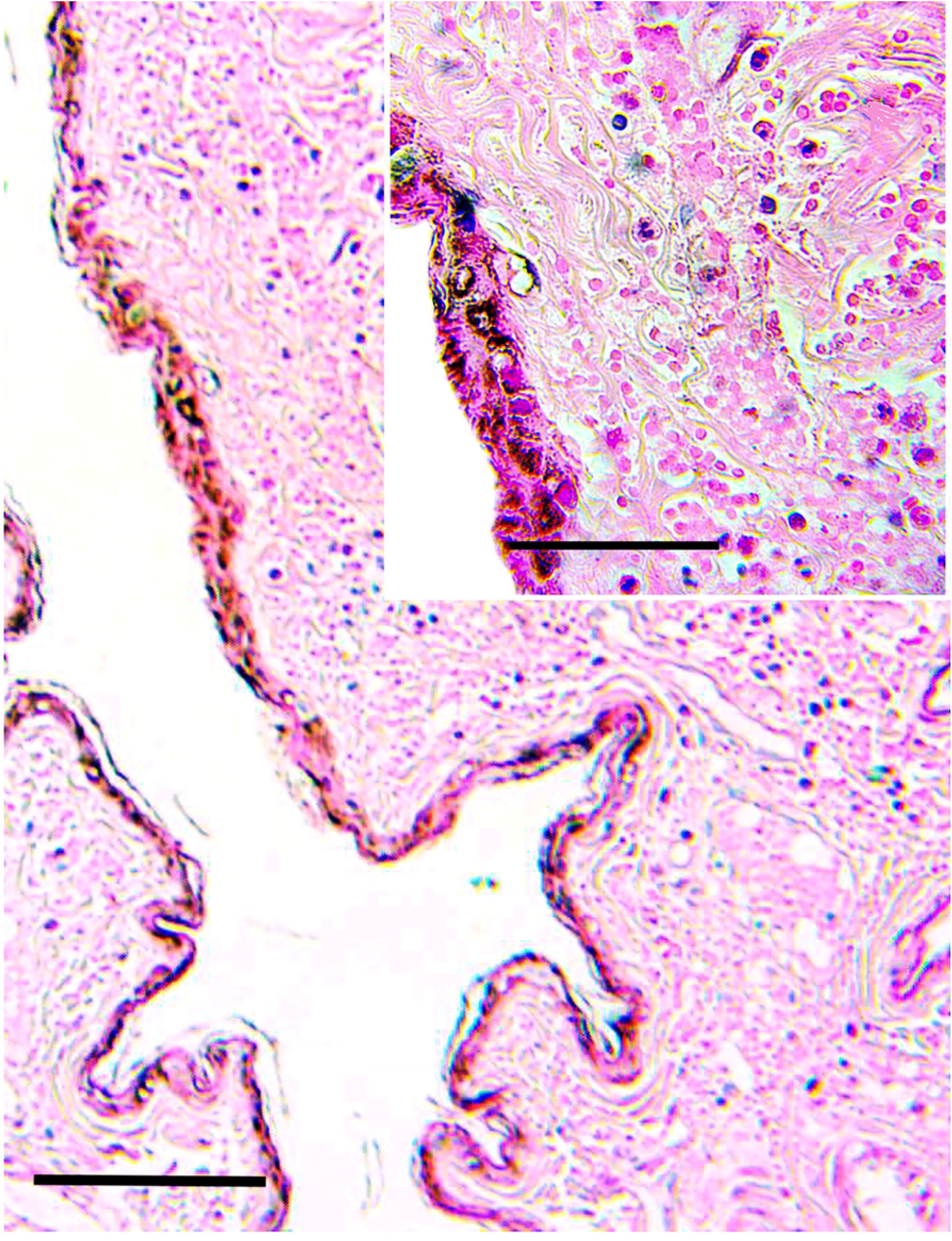
Photomicrographs of sections of the patagium. H&E stain; bar = 200 μm. The dermis has hemorrhages and has a mild interstitial neutrophilic infiltrate (*inset)*. H&E stain; bar = 50 μm.

**Figure 4.**
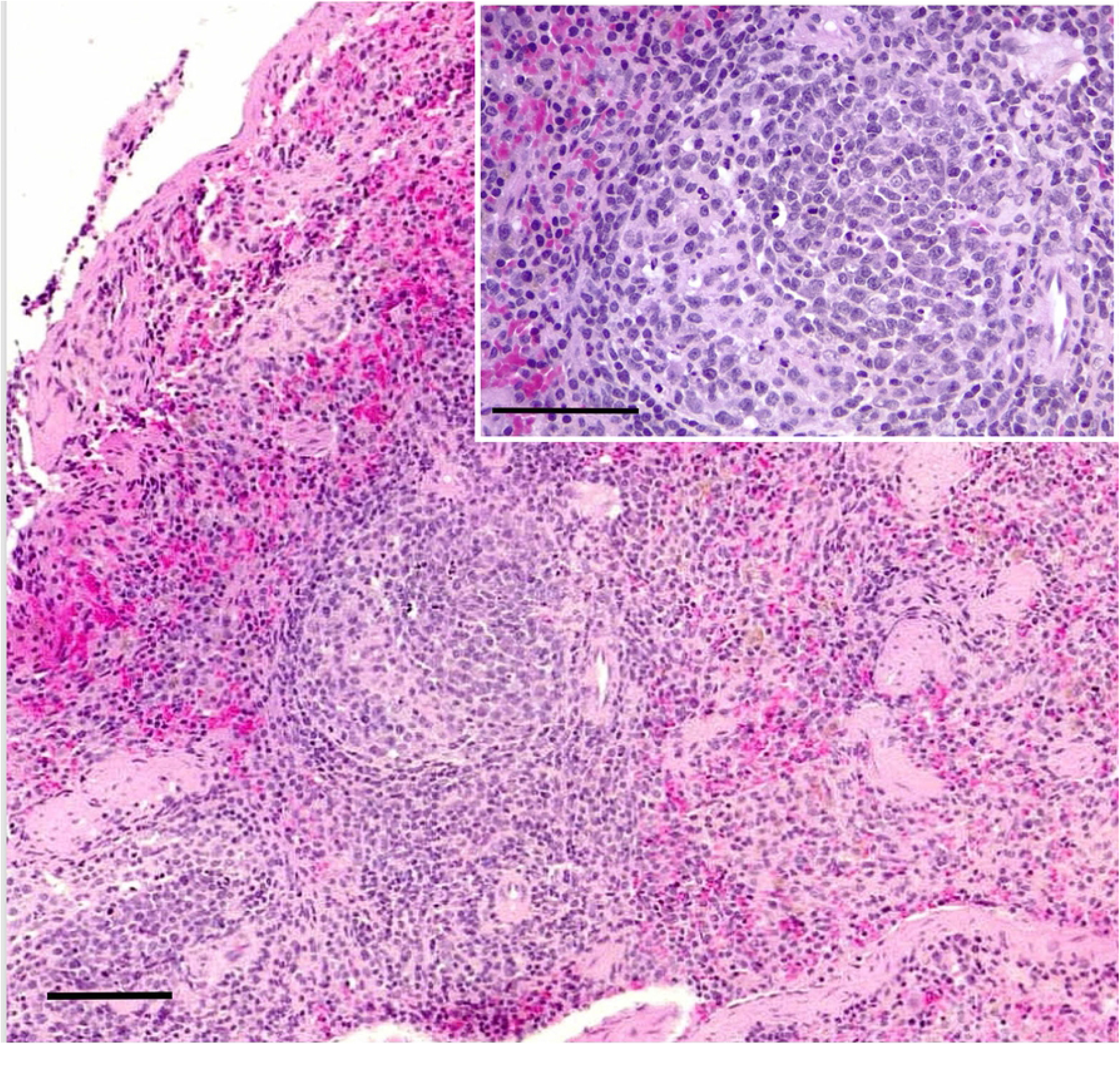
Photomicrographs of sections of the spleen. The white pulp has marked lymphoid follicular hyperplasia. H&E stain; bar = 100 μm. Lymphoid follicles have prominent germinal centers. H&E stain; bar = 50 μm.

#### Limited detection of viral RNA in urine

Consistent with Malmlov et al. 2019 (28), that detected ZIKV in low copy numbers (1×10^2^-1×10^3^ copies/ml) in the brain and urine in three individual bats, we detected ZIKV in only two urine samples of male individuals (20% detection rate, CI:4.5-52.0%) in urine from bat MO08 in 3 d.p.i. and MO05 in 9 d.p.i ranging 3.19 ×10^2^-3.61 ×10^2^ copies/ml (Table 1 A). All other animals and tissues tested negative.

#### Seroneutralization test

No serum had a neutralizing effect on the ZIKV strain H/PF/2013, contrasting with Malmlov et al. 2019 (28) findings, that reported modest seroconversion by an ELISA test (3200 vs >12000 in convalescent human sera) in three bats at day 28 pi. Putative reasons for the contrasting findings in our bats include the higher specificity of PRNT versus ELISA. These results are consistent with the negative detection of ZIKV antibodies by PRNT in 7 African bat species (41). But are in contrast with the frequent detection of ZIKV antibodies in free ranging bats in African early studies (25, 27). A major weakness these early studies is the possible lack of differentiation amongst Flaviviruses due to the testing method (HIT).

### Latin American-wide bat sampling

#### No ZIKV detection on free-living bats in different countries of Latin America

A total of 2056 bats were sampled (blood for ZIKV detection by qRT-PCR) belonging to different neotropical species and countries (Peru, French Guyana, and Costa Rica). In French Guyana, samples were collected over the years 2010, 2011, 2012, 2015, 2016, and 2017; in Costa Rica over 2017, 2018, and 2019; and in Peru, samples were collected only over 2017 and 2018. Sampling included 6 bat families and 33 genera of which 22 were family Phyllostomidae (*Anoura, Artibeus, Carollia, Chiroderma, Dermanura, Desmodus, Glossophaga, Linchonycteris, Lionycteris, Lonchophylla, Lonchorhina, Lophostoma, Mimon, Mycronycteris, Phylloderma, Phyllostomus, Platyrrhinus, Rhinophylla, Sturnira, Tonatia, Trachops*, and *Uroderma*); 3 were family Molossidae (*Cynomops, Eumops* and *Molossus*); 3 where Vespertilionidae (*Eptesicus, Myotis* and *Rhogeessa*); 3 were family Emballonuridae (*Peropteryx, Rhynchonycteris*, and *Saccopteryx*); one was family Noctilionidae (*Noctilio*), and one was family Mormoopidae (*Pteronotus*) within these families, 60 species were identified (Table S1). The largest number of sampled individuals belonged to the genus *Pteronotus* sp. (n=470, 23%), followed by the species *Carollia* sp. (n= 347, 17%); *Anoura* sp. (n=286, 14%), and *Molossus* sp. (n=282 13.7 %). No blood sample resulted positive to ZIKV by qPCR. This lack of detection is consistent with a serosurvey of Brazilian and African bats that also reported a lack of detection (41, 42). However, a sampling bias cannot be excluded, due to the diversity of bat species in Latin America. Urine samples were not analyzed and it must be considered that some studies in humans show that ZIKV RNA can be detected at higher levels and for a longer time after the onset of infection in the urine compared to blood and other fluids (29, 43-45).

## DISCUSSION

Our findings confirm and expand 28 Malmlov et al.’s (28) findings in 9 captured *Artibeus lituratus* bats from both sexes. Only two out of 180 (1%) urine samples, taken from ZIKV inoculated animals, showed low ZIKV copy numbers by qPCR. All other urine samples swabs and tissues from inoculated animals resulted negative. These results together with the lack of seroconversion founded suggest that *Artibeus* bats are not efficient amplifiers or reservoirs of ZIKV, such as what is observed for DENV (22, 23). In Malmlov’s work (28), only 3 positive samples by PCR were reported, two of them were urine samples and the third was nervous tissue, in the present work only two samples resulted positive and both were urine samples, in addition, some studies in humans show that ZIKV RNA can be detected at higher levels and for a longer time after the onset of infection in the urine compared to blood and other samples (43-45); thus, urine may be used as a preferred sample for further studies aiming to detect ZIKV in wild bats.

Histopathological findings are consistent with experimental infections in male *Artibeus* bats (28). In addition, the results are reminiscent of experimental murine ZIKV infections, which reported severe infection with inflammatory infiltration and degeneration of reproductive tissues and cells, but in these cases infections were performed by genital route in mice infected during diestrus-like phase (46) or by parenteral route in transgenic or treated mice lacking components of the innate antiviral response (47, 48)

The genital alterations observed suggest bat reproduction might be to some level affected by ZIKV. Further studies analyzing if the degeneration observed leads to infertility or alterations during pregnancy will be needed to assess if ZIKV could causes a disruption in bat reproductive health. Nevertheless, in the Americas, Zika has been closely associated with *Aedes* genus mosquitoes (2), especially *Ae. aegypti*, which are highly anthropophilic. Therefore, ZIKV would hardly reach wild bats.

Between days 3 and 7 after infection, clear leukopenia was observed in infected animals with subsequent recovery after day 9 pi. Generally, in mammals, as the first line of defense against infectious processes, the immune reaction includes leukocytosis in association with fever and during viral infections they can also occur with leukopenia and subsequent lymphocytosis as happened in our research. However, in some lipopolysaccharide-stimulated insectivorous and neotropical bats (LPS), no leukocytosis or fever was observed [41, 42]. Another feature of leukocyte kinetics reported in Vespetilionidae bats in Karelia is the clear leukopenia generated during hibernation in bats [43]. These authors conclude that the differences in the proportion of lymphocytes may be related to the biological, ecological and physiological peculiarities of the bats studied and generates the need for further studies to determine a correlation between bat infection with ZIKV and leukocyte kinetics of *Artibeus*.

In summary, neotropical bats from different families might not have an important role in ZIKV transmission dynamics.

## Ethical statements

Capture and animal handling was approved by the Mexican environmental standards (permit SEMARNAT Mexico No SGPA/DGVS/08986/18).

The study and associated protocols were design based on national ethical legislative rules and approved by Institutional Committee of Care and Use of Animals of the University of Costa Rica (CICUA-042-17), Committee of Biodiversity of the University of Costa Rica (VI-2994-2017), National System of Conservation Areas (SINAC): Tempisque Conservation Area (Oficio-ACT-PIM-070-17), La Amistad-Caribe Conservation Area (M-PC-SINAC-PNI-ACLAC-047-2018). Authorization for bat capture in French Guiana was provided by the Ministry of Ecology, Environment, and Sustainable Development during 2015–2020 (approval no. C692660703 from the Departmental Direction of Population Protection (DDPP, Rhône, France). All methods (capture and animal handling) were approved by the Muséum national d’Histoire naturelle, Société Française pour l’étude et la Protection des Mammifères, and the Direction de l’environnement, de l’aménagement et du logement (DEAL), Guyane. Bat sampling in Peru was authorized by the Research Ethics Committee of the Institute of Tropical Medicine “Daniel Alcides Carrion” (Constancia de aprobación CIEI-2017-16)

The survey did not involve endangered or protected species

## Acknowledgments

Zika virus was kindly donated by Dr. Luis Antonio Alonso Palomares and Dra Isabel Salazar from the Virology and Immuno-virology Laboratory of the “Escuela Nacional de CienciasBiológicas del Instituto Politécnico Nacional, Mexico”, ENCB-IPN, Mexico City. This work was supported by the German Centre for Infection Research (DZIF) through the ZIKApath project, and the European Union’s Horizon 2020 research and innovation program through the ZIKAlliance project (grant agreement 734548). Mexican work was partially supported by CONACYT – FONCICYT Project “Una Alianza Global para Controlar y Prevenir el Virus del Zika” Number 274386. Costa Rica work was also supported by FEES-CONARE B7362 project.

AMS was funded by the German Academic Exchange Service

(DAAD) and The Office of International Affairs and External Cooperation (OAICE) from the University of Costa Rica. We wish to acknowledge Juan Enrique del Águila Romero, Executive Director of the Red de Salud de Alto Amazonas, his logistical support in the field work, and Dr. Vilma R. Béjar Castillo, Director of Instituto de Medicina Tropical “DAC” his support to the project. Also, thanks Kike Sinarahua Ishuiza, Estefany Janneth Garay Vela and Antolín Saldaño from Salud Ambiental de la Red de Salud de Alto Amazonas their contribution to field works.

## Figure legends

**Table S1**. List of bats sampled in French Guiana, Peru and Costa Rica from 2010 to 2017 for the detection of ZIKV in blood by rtPCR and antibodies by seroneutralization test.

**Figure S1**.- Recorded corporal temperature of control and inoculated ZIKV bats, from 0 to 21 days pi.

Minimum temperature: 34.1 º C

Maximum temperature: 38.5 º C

Geometric mean of inoculated bats: 35.6 ºC; SD 0.75; CI 95%

Geometric mean of control bats: 35.7 ºC; SD 0.6; CI 95%

